# Throat colour polymorphism in relation to sex and body size of the Litter skink, *Lankascincus fallax*

**DOI:** 10.1101/2022.10.18.512657

**Authors:** J.M.A. Ishara K. Jayamanna, Anslem de Silva, Kanishka D.B. Ukuwela

## Abstract

Colour polymorphism is a pervasive phenomenon in both animal and plant kingdoms and understanding its evolution and maintenance is of great interest. Among the lizards of Sri Lanka, the endemic skink *Lankascincus fallax* shows throat colour polymorphism in which, the underlying basis is not clearly known. In this study, we examined the relationship of the three different throat colour morphs observed in this species with the sex, body size and the geographic location of *L. fallax*. Live skinks were sampled from two locations in Sri Lanka and sex and the throat colour was categorized visually and the snout to vent length (SVL) was measured. Tail tips of some selected individuals from the two locations were taken and a fragment of the 12S rRNA gene was sequenced in representative individuals having the different throat colour morphs. Pairwise genetic distance of the three colour morphs ranged between 0.4–0.5% confirming that the three colour morphs were the same species. Three colour morphs (red, black and white) were observed in males in both locations, while only the white morph was observed in females, suggesting that the throat color polymorphism was confined to males. There was a significant difference between the mean SVL of males with red and black throat colours (39.35 mm) and males with white throat colours (30.31 mm). Thus, the study suggests that the throat colour in these skinks is highly associated with sex and the body size in males. The study further suggests that *L. fallax* is sexually dichromatic and that the males show throat colour polymorphism. However, future studies are necessary to understand the underlying drivers for the presence and maintenance of sexual dichromatism and throat colour polymorphism in *L. fallax*.

## INTRODUCTION

Colour polymorphism of an organism is the display of several heritable colour variants in individuals of the same age and same sex in a population for which the expression is sensitive neither to the environment nor to the body condition (Cooke & Buckley, 1987). Processes such as genetic drift and gene flow, disruptive selection, heterosis, apostatic selection, sexual selection and sensory bias help in the development and maintenance of colour polymorphism in species (Galeotti et al., 2013). Colour polymorphism helps to represent the physiological or fitness characters of polymorphic species and accomplish functions, such as status of sexual signaling, crypsis, mutualism, aposematism, mimicry, thermoregulation and habitat use (Galeotti et al., 2013). Further, it may result in variances among morphs in morphological, physiological, life history or behavioral traits, such as body size, clutch size, immune function, and antipredatory behavior, that may initiate the divergence of species (B Sinervo & Svensson, 2002; Calsbeek et al., 2010). However, the risk of extinction of polymorphic species as compared to non-polymorphic species is less (Ducatez et al., 2017). Thus, colour polymorphism may perform as a buffer against drastic environmental changes (Ducatez et al., 2017) and assit to establish more successfully in new areas and is considered to be less vulnerable to extinction indicating that colour polymorphism promotes species survival (Forsman et al., 2012).

Genetically inherited colour polymorphism is found in many species throughout the animal and plant kingdoms (Roulin, 2004). Some well-known examples of colour polymorphism among reptiles are found among snakes (K. D. B. Ukuwela & Dawundasekara, 2012; Pizzatto & Dubey, 2012) and lizards of the families, scincidae (Chapple *et al*., 2008), lacertidae (Huyghe et al., 2009), agamidae (Teasdale & Stevens, 2013) and geckoes (Ota et al., 1995; Bauer & Branch, 2012; Flecks et al., 2012; Rösler et al., 2012). Despite the drivers of colour pattern variation, body colouration and patterns are not reliable investigative characters for reptile taxonomy, because they can considerably be dissimilar between or even within populations, between sexes, with ontogeny, activity, physiological state, time of day and parasite load (Mats Olsson et al., 2013). Thus, in reptile taxonomy and determination of species status, colour polymorphism plays a vital role. Among reptiles, lizards often express a high intraspecific variability of colour patterns and therefore are an excellent model to study the evolution and maintenance of colour polymorphism (Galeotti et al., 2013).Therefore, investigation of colour polymorphism is important to understand evolution of life history in lizards.

Among the lizards of Sri Lanka, the endemic skink species *Lankascincus fallax* shows throat colour polymorphism. The *Lankascincus* genus which is endemic to Sri Lanka comprises 10 species of small (10–60 mm SVL) litter dwelling skinks (Greer, 1991; Batuwita, 2019). *Lankascincus fallax* is the most abundant and widely distributed of all *Lankascincus* species in Sri Lanka being distributed in all the bioclimatic zones from the coast to the Central Hills except elevations above 1,200 m in Sri Lanka (Somaweera, R., 2009; De Silva & Ukuwela, 2017). Most preferable habitats of this species are leaf litter, under stones, decaying logs, and within debris (Batuwita, 2019). High moisture level in leaf litter is crucial for the existence of these species (Batuwita, 2019). Three different types of throat colour morphs (white, red with white spots and black with white spots) are shown by *L. fallax* (Fig. 1). The morph with red coloured throat was initially assigned to a different species, *Sphenomorphus rufugulus* (=*Lankascincus rufugulus*) (Taylor, 1950; Goonatilake et al., 1999; Balasubramaniam & Krishnarajah, 2004). However, due to lack of other morphological differences, *S. rufugulus* was later synonymized with *L. fallax* (Das & De Silva, 2005; Wickramasinghe *et al*., 2007). It is speculated that the throat colour polymorphism in this species is related to the sex and the maturity of the lizard (Wickramasinghe et al., 2007; Batuwita, 2019). However, these speculations are yet to be tested. In this study we examined whether the three different throat colour morphs belongs to same species or not through DNA barcoding. As our analyses indicated that the three colour morphs were the same species (see below), we then examined whether there are relationships between throat colour and body size, sex and the geographic location.

**Figure 1.**
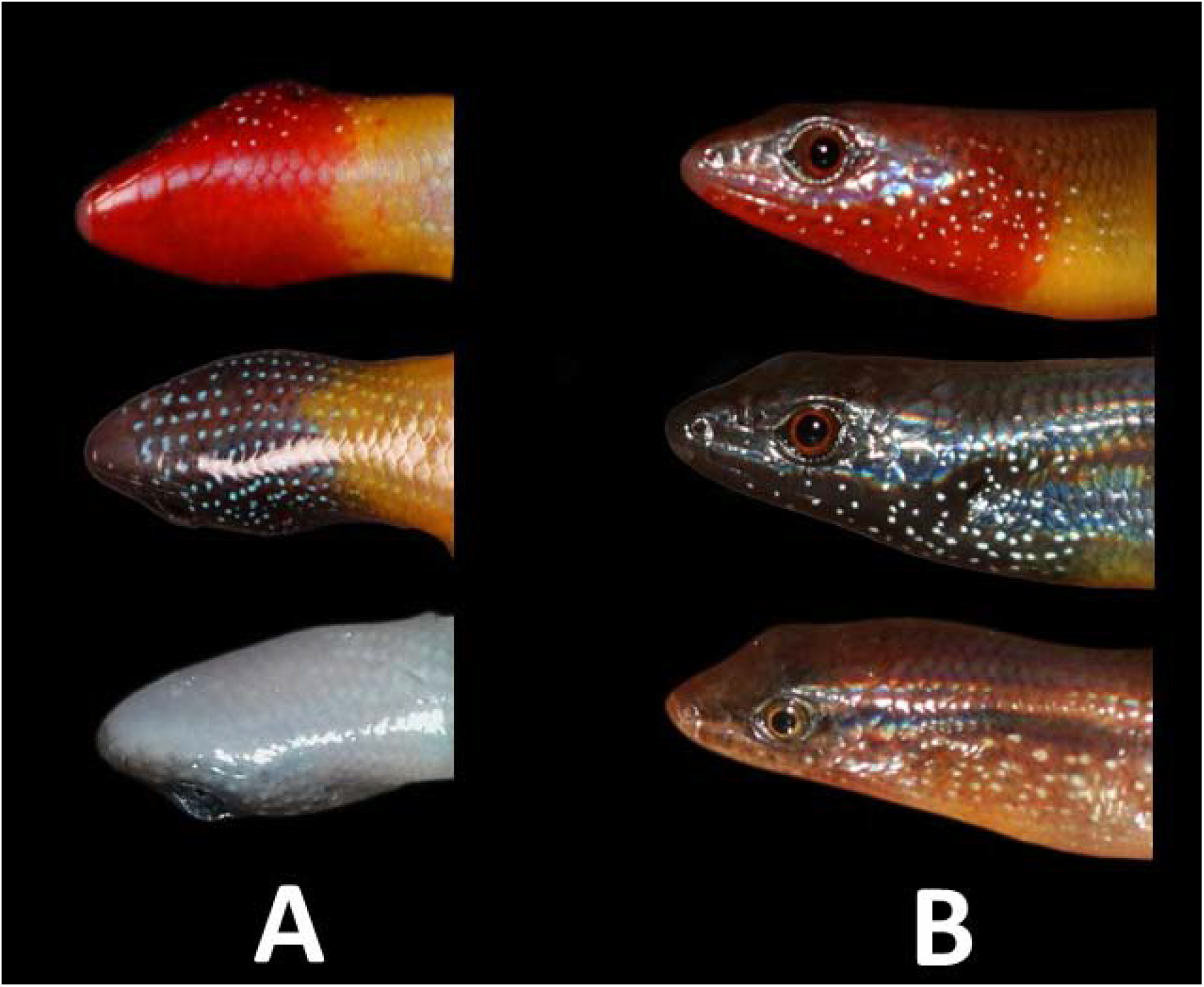
Three colour morphs of *Lankascincus fallax*. (a) Ventral side of throats of three different throat colour individuals, (b) Lateral view of throats of three different throat colour individuals

## MATERIALS and METHODS

Individuals of *L. fallax* was sampled from two locations (Mihintale and Polgahawela) in Sri Lanka. These two locations (112 km apart) were selected to represent two bioclimatic zones in Sri Lanka, the Dry zone (annual rainfall between 1200–1800 mm) and the Intermediate zone (annual rainfall between 1800–2500 mm. Mihintale (8.3540, 80.5035, Anuradhapura district, Northcentral province, Sri Lanka) belongs to the dry bioclimatic zone of Sri Lanka while Polgahawela (7.3576, 80.3429, Kurunegala district, Northwestern province, Sri Lanka) belongs to the intermediate bioclimatic zone of Sri Lanka. These two sites were selected in order to compare the inter-population variation in the throat colours representing the two bioclimatic zones. The habitats of both locations were highly similar having mostly anthropogenic habitats including home gardens with leaf litter, grasslands and plantations with coconut and banana.

The study was carried out by catching live skinks from November 2018 to February 2019 to determine throat colour polymorphism through external observations. A 3mm portion of the tail tips were taken from three individuals from each colour morph from each sampling site for DNA analyses to confirm the identity of the species. These tail tips were preserved in 1.5 ml microcentrifuge tubes containing 90% ethanol. Taking tissue samples from tails does not harm skinks as they usually lose their tails to avoid predation and eventually regrow the tail. Sampling was done once a month from each population in Mihintale and Polghawela. More than 30 individuals were captured by hand during the daytime through active searching within the study area in each visit. All captured individuals were put into plastic containers until throat colour and sex were determined, and the measurements were taken. The date of sampling, time, habitat and location were noted each time.

In each individual, head length, head width, snout to vent length and tail length were measured. All measurements were taken using a Vernier caliper to the nearest 0.02 mm. Sex of these skinks was determined by hemi-penal trans illumination technique (Brown, 2009). Throat color was observed through normal eyesight and was assigned to either of the three throat colour morphs; red with white spots, black with white spots and white. The number of eggs in females were observed by external examination (Ukuwela, 2009). After all the measurements and observations were made, the skinks were released back to the same collection site.

To confirm that different colour morphs of the skinks belonging to the same species, mitochondrial 12S rRNA gene was sequenced and examined. Mitochondrial 12S rRNA gene was used as this gene is widely used in species identification and phylogeny estimation of vertebrates because of its highly conserved priming sites and high percentage of nucleotide substitutions among closely related species (Kocher et al., 1989; Yang et al., 2014). Whole genomic DNA were extracted from the tail tissue samples representing the red, black and white throat colored individuals using Promega Wizard**®** genomic DNA purification kit (Promega Corporation, Madison, Wisconsin, USA) following manufacturer’s protocols. DNA extracts were quantified using a Qubit Fluorometer (ThermoFisher Scientific, Waltham, Massachusetts, USA) and extracts greater than 10 ng/μlwere used in the Polymerase Chain Reaction (PCR) amplification. 12S rRNA gene regions were PCR amplified using primers AAACTGGGATTAGATACCCCACTAT and GAGGGTGACGGGCGGTGTGT (Kocher et al., 1989). The PCR amplification was carried out in 25 μl volumes employing 35 cycles with an annealing temperature of 55 °C following standard PCR protocols with Promega PCR master mix (Promega Corporation, Madison, Wisconsin, USA). The success of the PCR amplification of each sample was confirmed through Gel electrophoresis using 5 μl of each sample. The successfully amplified PCR products were then sequenced in both directions at Genetech Sri Lanka Pvt. Ltd, Colombo. Consensus sequences from forward and reverse reads were assembled in GENEIOUS PRO 5.6 software. DNA sequences generated in this study are accessioned in the NCBI GenBank under the accession numbers, #######–#######. All the consensus sequences were aligned in GENEIOUS PRO 5.6 and genetic distances among the sequences were calculated to determine the amount of genetic divergence between DNA sequences of individuals of different colour morphs. A distance of 2% was used as a cut off value to determine whether the individuals of different colours belonged to the same species or not (Johns & Avise, 1998).

The relationships between sex versus throat colour was examined using Pearson’s Chi-squared test. Relationship between body size (SVL) and the throat colour of males was examined using Kruskal-Wallis rank-sum test. A Pearson’s Chi-squared was used to examine the relationship between throat colour and the sampling time of the year. All statistical tests were done using R statistical software 3.5.2 (R development core team, 2013).

## RESULTS

### DNA Analysis

The average corrected (HKY) pair-wise genetic distance between the 12S rRNA gene sequences generated from the tissue samples representing the three colour morphs ranged between 0.4–0.5% (Table 1). Since, these values are lower than the 2% cutoff levels of species delimitation for mitochondrial genes (Johns & Avise, 1998) the three colour morphs therefore most likely belong to the same species.

**Table 1.** Pairwise corrected (HKY) genetic distances (%) between three different colour morphs of *Lankascincusfallax*

|  | Black (n=2) | Red (n=2) | White (n=2) |
| --- | --- | --- | --- |
| Black | - |  |  |
| Red | 0.5 | - |  |
| White | 0.5 | 0.4 | - |

### Relationship between the throat colour and sex

All the sampled males in both locations were represented by three throat colours (red with white spots (Red), black with white spots (black) and white) however all the females only had white coloured throats in all four different months (Table 2).

**Table 2.** Sex and throat colours of different individuals of *Lankascincus fallax* observed during the study period from Polgahawela and Mihintale.

|  | November |  | December |  | January |  | February |  |
| --- | --- | --- | --- | --- | --- | --- | --- | --- |
|  | Females | Males | Females | Males | Females | Males | Females | Males |
| Red | 0 | 11 | 0 | 3 | 0 | 4 | 0 | 12 |
| Black | 0 | 8 | 0 | 13 | 0 | 8 | 0 | 13 |
| White | 35 | 3 | 49 | 11 | 32 | 9 | 41 | 18 |
| Total | 35 | 22 | 49 | 27 | 32 | 21 | 41 | 43 |

Pearson’s Chi-squared test indicated highly significant (χ^*2*^ **=** 134.01, df = 1, p≤0.000) association between sex and the throat colour (coloured versus non colour throated individuals) of *L. fallax*. Similarly, a Pearson’s Chi-squared test indicated highly significant association (χ^*2*^ **=** 134.01, df = 2, p≤0.000) between sex and throat colour in *L. fallax* (three throat colours Red, black, white).

### Relationship between the throat colour of males and body size (SVL)

The mean SVL of red and black-throated males was 39.36 mm (SD = ±2.31) while it was 31.93 mm (SD = ±7.55) in white-throated males. A Welch’s Two Sample t-test indicated that there is a significant difference between mean SVL of colour throated (red and black) and white-throated males (t = 11.458, df = 51.543, p≤0.000). This indicates, that SVL of red and black-throated males was greater than SVL of non-coloured males (Fig. 2). Among the red and black-throated males, black-throated males had slightly larger mean SVL (39.72 mm (SD=±2.28)) compared to red-throated males (mean SVL = 38.83mm (SD=±2.28) mm) (Fig. 3). However, Welch’s Two Sample t-test indicated that there is no significant difference between mean SVL of red-throated and black-throated males (t = 1.616, df = 60.876, p=0.112).

**Figure 2.**
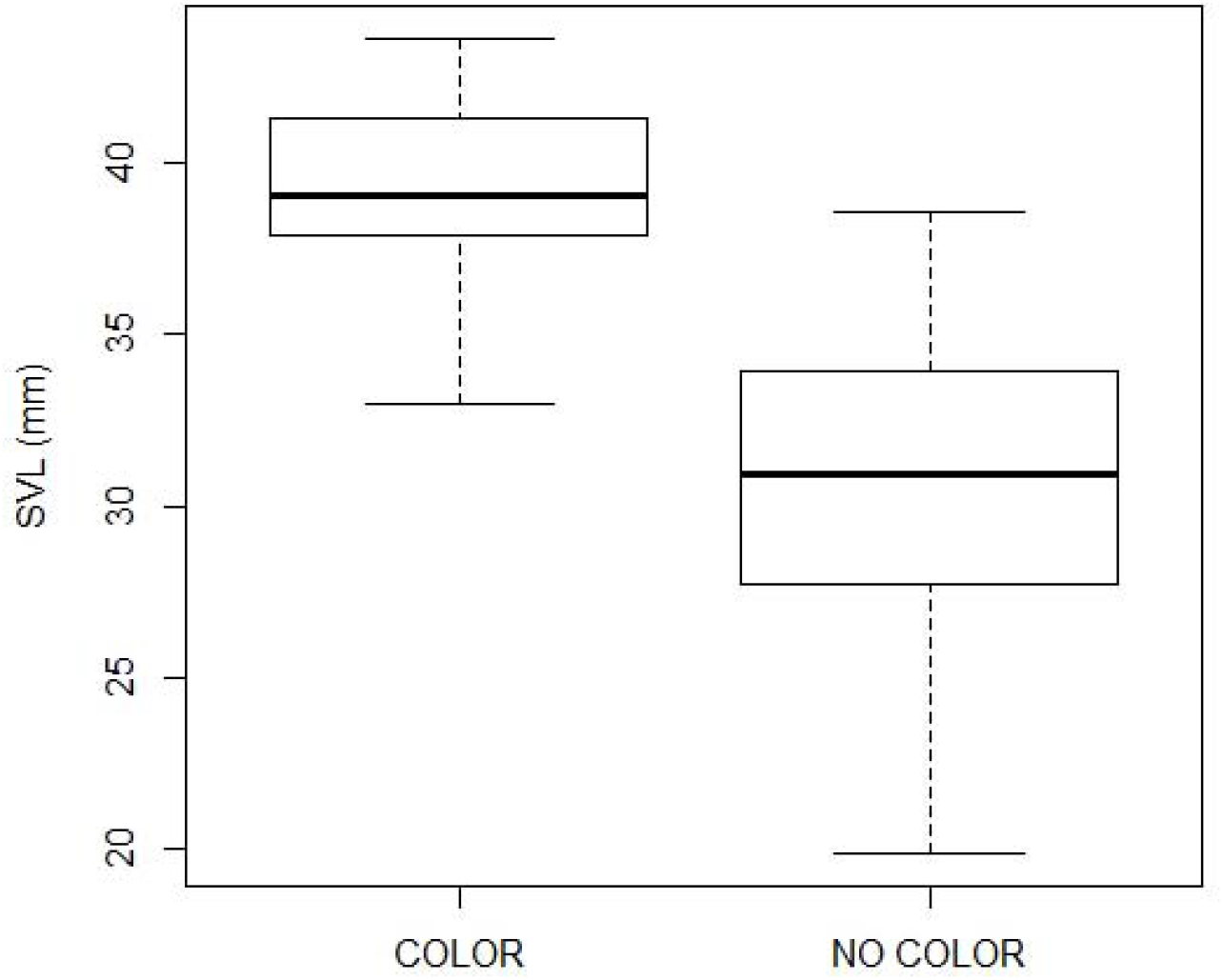
Mean SVL of coloured and non coloured male individuals sampled.

**Figure 3.**
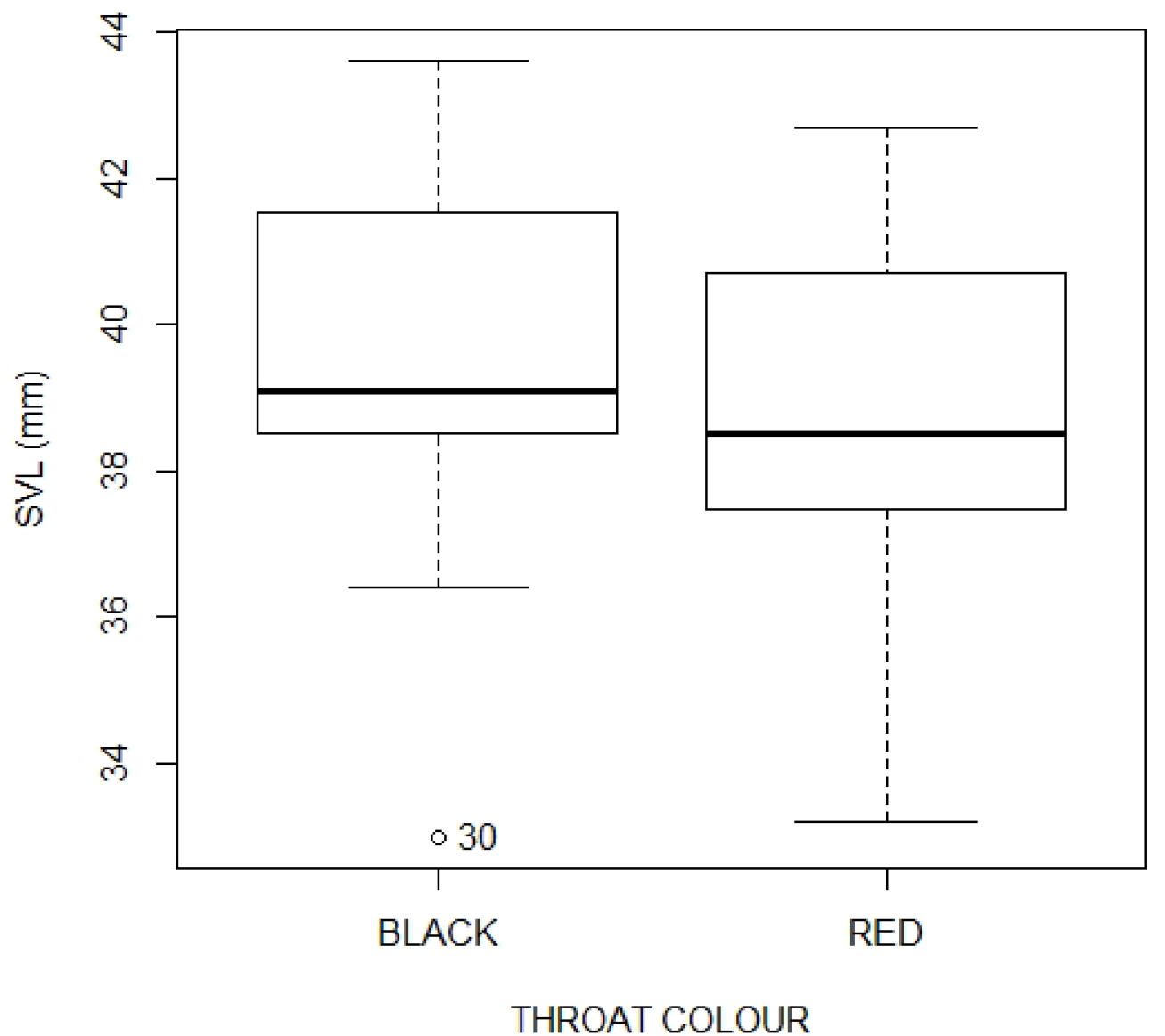
Mean SVL distribution of black and red throated male individuals sampled.

**Figure 4.**
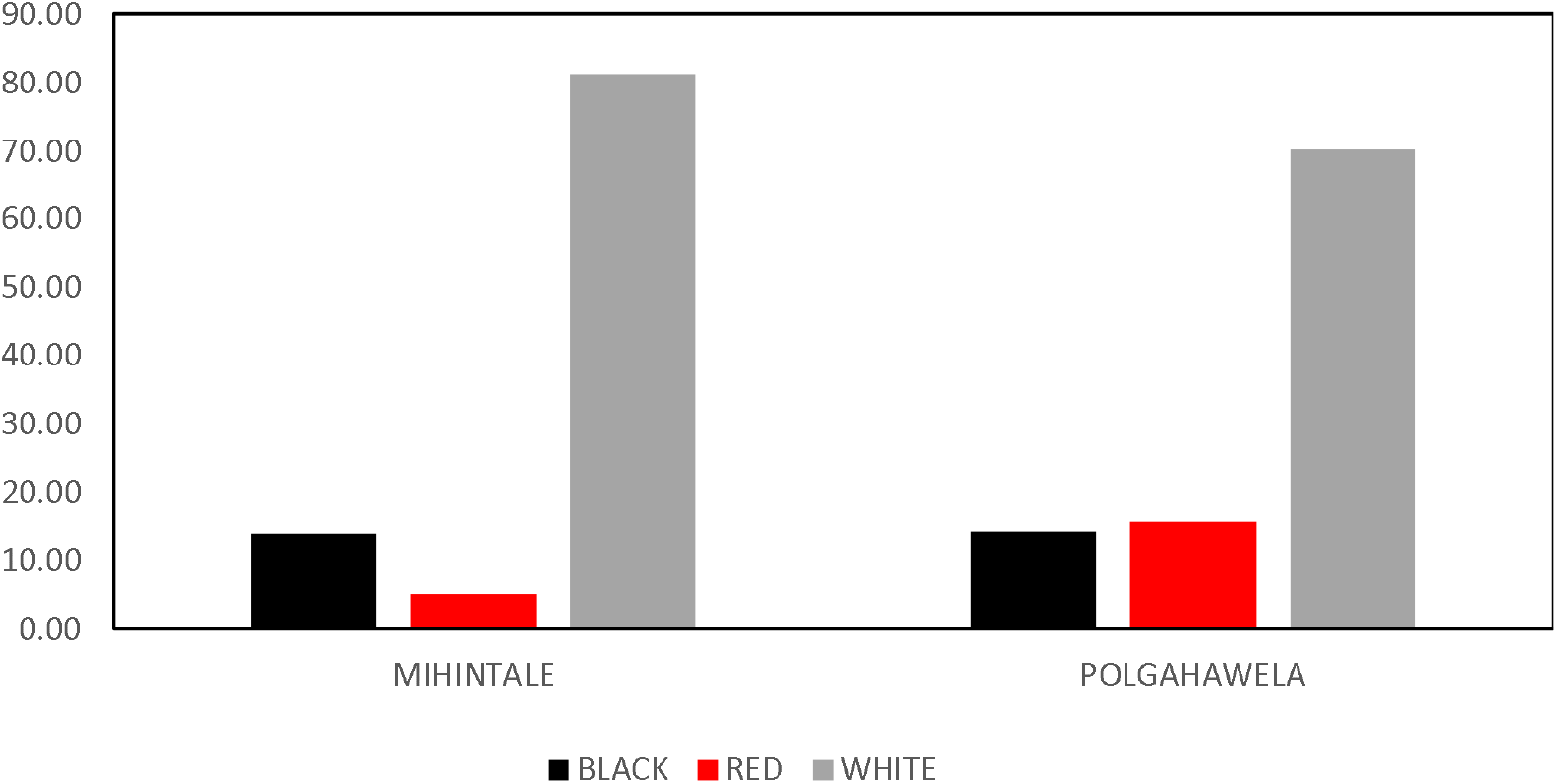
Throat colour percentages of *Lankascincus fallax* in the two sampling sites.

### Relationship between throat colour and the sampling location

The percentages of the number of individuals belonging to the three throat colours were different between the two locations (Table 3). This difference was highest among the percentage of individuals having red coloured throats between the two locations (i.e. Polgahawela = 15.71%, Mihintale = 5.03%). A Pearson’s Chi-squared test indicated that there was a significant difference (χ^*2*^= 9.694, df = 2, p-value = 0.007) between the numbers of individuals having different throat colours at the two locations.

**Table 3.** Number of individuals with different throat colours observed at the two sampling locations. The percentage of each category to the total number of individuals observed is given in parenthesis.

| Location | Black | Red | White |
| --- | --- | --- | --- |
| Mihintale | 22 (13.83%) | 08 (5.03%) | 129 (81.13%) |
| Polgahawela | 20 (14.28%) | 22 (15.71%) | 98 (70.00%) |

## DISCUSSION

Our findings show that red and black throat colouration is restricted to males while white throat colouration is present in both males and females of *L. fallax*. The study further revealed that the males with red and black throats are longer than males with white throats and that the frequency of throat colours between the two locations is different. We discuss these findings with respect to sexual dichromatism, colour polymorphism and geographic variation in colour polymorphism.

Male *L. fallax* from both locations were represented by three throat colours (red, black and white) while all the females only had white coloured throats indicating that distinctly bright throat colouration is restricted to males. This phenomenon which is known as sexual dichromatism where one sex is distinctly or brightly coloured than the other is widespread in the animal kingdom and is well known from fish (Kingston et al., 2003), lizards (Barry Sinervo & Lively, 1996; M Olsson et al., 2007; Sacchi et al., 2013; James E Paterson & Blouin-demers, 2017) and birds (Burns & Shultz, 2012). One of the most well known underlying causes for sexual dichromatism is female mate preference to certain colour morphs (Stuart-Fox & Ord, 2004; Chen et al., 2012; Rossi et al., 2019). However, *L. fallax* is not only sexually dichromatic, the males also show throat colour polymorphism. Male colour polymorphism is also known to be strongly influenced by sexual selection (Healey et al., 2007; Wellenreuther et al., 2014). Previous studies have shown that females show a higher preference to a particular male colour morph when there are multiple colour morphs among males in the population (Kingston et al., 2003; Stuart-Fox & Ord, 2004; Healey et al., 2007; Rossi et al., 2019). Interestingly, it has also been shown that male throat colour polymorphism is related to habitat selection in tree lizards (J. E. Paterson & Blouin-Demers, 2018). However, certain studies have found that the frequency of male colour morphs oscillate over time leading to the maintenance of different colour morphs in the populations (Barry Sinervo & Lively, 1996).

Contrastingly, having highly conspicuous colours are also known to increase the risk of predation (Stuart-Fox et al., 2003). These previous studies suggest that male colour polymorphism is strongly linked to mate choice and polymorphism is maintained in the population through frequency dependent selection. However, the drivers and maintenance of throat colour polymorphism in *L. fallax* are yet to be understood. Given its diminutive nature, observing mating behavior is challenging. Hence, nothing is known about the female mate preferences of *L. fallax*. Thus, future studies should be focused to examine the female mate preferences on throat colour morphs in *L. fallax* and how mate preferences change through time.

The larger SVL of males with red and black throats to that of males with white throats may indicate a higher level of maturity in males with brightly coloured throats. Although the males with black throats had a slightly larger SVL than males with red throats, this was not significant. Usually, larger body size (SVL) is an indication of a higher level of maturity or even sexual maturity. Thus, the size difference in males with respect to throat colours might indicate that body colour is closely related with the state of maturity. Previous studies indicate that the colour development may be a result of sexual maturity (Cox et al., 2005; Calisi & Hews, 2007). In males of Drakensberg crag lizard *(Pseudocordylus melanotus)*, the development of bright colours corresponds due to gonadal maturation (Mounton & Wyk, 1993). In *L. fallax*, it is most possible that throat colours (red, black) are developed in sexually mature males as the males with throat colours had a larger body size. If this is so, throat colour may be have a close link to female mate choice/preference where a certain throat colour may be preferred by females. However, these speculations should be confirmed through further studies. Nonetheless, our study cannot confirm any of these speculations as sexual maturity were not determined in these skinks due to the absence of external determinants.

The significant difference in throat colours between the two geographic locations indicates that there are differences in the frequencies of throat colours in the two populations. The percentage of males with red throats were almost three fold in Polgahawela compared to Mihintale. A previous study indicated that the proportion of black-throated males were greater compared to read throated males in Mihinthale (Kumari et al. 2010). However, in Polgahawela, the proportion of red-throated males was greater than black-throated males. The geographic variation in colour polymorphism has been observed in numerous species of animals including invertebrates (Grant et al. 10998, Ozgo 1998) and vertebrates (Galecotti et al. 2003; Corl et al. 2010, Mclean et al. 2014, Mclean and Stuart-Fox, 2014). Negative frequency dependent selection, secondary contact and hybridization of previously isolated populations, spatial variation in selection due to local adaptations and stochastic processes such as genetic drift and founder effects have been attributed to geographic variation in animal colour morphs (Mclean and Stuart-Fox, 2014). Due to sampling being limited to two locations, geographic variation of throat colour in *L. fallax* cannot be confirmed. However, a more broad scale sampling spanning the entire area of distribution is necessary to ascertain the prevalence of throat colour polymorphism in *L. fallax*.

The difference in the frequency of red-throated males between the two locations could also be due to seasonal variation as the two locations are situated in two distinct bio-climatic regions of Sri Lanka, which experience different rainfall seasonality. Seasonal variation of male colouration has been observed in many species lizards both in the tropical (Pal et al., 2011; Deodhar & Isvaran, 2017) and temperate regions (Germano and Williams 2007) of the world. These variations in the body colouration have been accounted for reproductive seasonality (Germano & Williams, 2007; Deodhar & Isvaran, 2017; Ortega et al., 2019) and seasonal environmental changes (Pellitteri-Rosa et al., 2020). However, a long-term study spanning multiple geographic locations and climatic seasons is necessary to ascertain the influence of seasonality on the geographic differences in throat colour polymorphism in *L. fallax*.

## CONCLUSION

The study suggests that the throat colour in *L. fallax* is highly associated with sex and the body length (most likely sexual maturity) in males. The findings further indicate that there is a difference in throat colour frequencies between the two study locations. Thus, future studies are necessary to understand the underlying drivers for the evolution, adaptive significance and maintenance of these different throat colours in *L. fallax*.

## ACKNOWLEDGMENTS

We thank the Department of Wildlife Conservation of Sri Lanka for the research permit (WL/3/2/70/15) to carry out this study and the Rajarata University of Sri Lanka for financial aid (RJT/RP&HDC/2016/FOAS/R/03). We are grateful to Rajnish Vandercone for the encouragement and the advice given during the project. Our sincere thanks go to Buddhima Kalani, Akila Weerasinghe, Udara Kaushalya, Shamesh Wijenayake, Dhanushka Sampath, Nalin Madushanka, Chathura Nagahawathha, Isuru Supasan, Rakshitha Bandara and Ujith Rasanjana for the assistance given in the field.

## Notes

### Competing Interest Statement

The authors have declared no competing interest.

